# Somatosensory cortex microstimulation modulates primary motor and ventral premotor cortex neurons with extensive spatial convergence and divergence

**DOI:** 10.1101/2023.08.05.552025

**Authors:** Brandon Ruszala, Kevin A. Mazurek, Marc H. Schieber

**Affiliations:** Department of Biomedical Engineering, University of Rochester, Rochester, NY, 14627; Department of Neurology, University of Rochester, Rochester, NY, 14642; Department of Neuroscience, University of Rochester, Rochester, NY 14642

**Keywords:** Brain-computer interface, Brain stimulation, Instruction, Intracortical microstimulation, Excitation, Inhibition

## Abstract

Intracortical microstimulation (ICMS) is known to affect distant neurons transynaptically, yet the extent to which ICMS pulses delivered in one cortical area modulate neurons in other cortical areas remains largely unknown. Here we assessed how the individual pulses of multi-channel ICMS trains delivered in the upper extremity representation of the macaque primary somatosensory area (S1) modulate neuron firing in the primary motor cortex (M1) and in the ventral premotor cortex (PMv). S1-ICMS pulses modulated the majority of units recorded both in the M1 upper extremity representation and in PMv, producing more inhibition than excitation. Effects converged on individual neurons in both M1 and PMv from extensive S1 territories. Conversely, effects of ICMS delivered in a small region of S1 diverged to wide territories in both M1 and PMv. The effects of this direct modulation of M1 and PMv neurons produced by multi-electrode S1-ICMS like that used here may need to be taken into account by bidirectional brain-computer interfaces that decode intended movements from neural activity in these cortical motor areas.

**Significance Statement:** Although ICMS is known to produce effects transynaptically, relatively little is known about how ICMS in one cortical area affects neurons in other cortical areas. We show that the effects of multi-channel ICMS in a small patch of S1 diverge to affect neurons distributed widely in both M1 and PMv, and conversely, individual neurons in each of these areas can be affected by ICMS converging from much of the S1 upper extremity representation. Such direct effects of ICMS may complicate the decoding of motor intent from M1 or PMv when artificial sensation is delivered via S1-ICMS in bidirectional brain-computer interfaces.

## Introduction

Intracortical microstimulation (ICMS) injects brief microampere current pulses into cortical gray matter, eliciting action potentials in neural elements in the vicinity of the electrode tip. Early work found that single 40 µA pulses delivered in cat motorsensory cortex elicited action potentials in neuron somata up to 400 µm away (Stoney et al., 1968). Subsequent studies have demonstrated that ICMS pulses delivered in mouse and cat visual cortex also elicit action potentials in local axons of passage, producing antidromic action potentials in sparse populations of cortical neurons up to millimeters from the electrode tip (Histed et al., 2009). A recent computational model of macaque somatosensory cortex further suggests that all ICMS-elicited action potentials originate in axons, and that the radius of activated axons increases with increasing current through the typical range from 15 to 100 µA. Antidromic propagation of those elicited action potentials leads to an increasing density of antidromically activated somata over a distance up to ∼2 mm, and while less frequent, orthodromic propagation of those action potentials can elicit somatic action potentials transsynaptically, which may affect neurons at even further distances (Kumaravelu et al., 2022). Indeed, individual ICMS pulses delivered in macaque primary motor cortex (M1) or ventral premotor cortex (PMv) elicit action potentials in local corticospinal neurons transsynaptically (Maier et al., 2013).

However they originate, orthodromically conducting action potentials elicited by ICMS can produce transsynaptic effects in additional neurons at considerable distances. While individual ICMS pulses generally may be subthreshold for eliciting action potentials transsynaptically, such effects become prominent when ICMS pulses are delivered in high-frequency trains (≥ 200 Hz), enabling temporal summation of successive excitatory postsynaptic potentials (Jankowska et al., 1975; Hussin et al., 2015). Trains of ICMS pulses delivered in motor cortex excite distant spinal motoneurons and produce muscle contractions (Asanuma and Ward, 1971; Andersen et al., 1975). Delivered at lower frequencies in the primary somatosensory cortex (S1), ICMS pulse trains elicit somatosensory percepts (Romo et al., 1998; Flesher et al., 2016; Callier et al., 2020).

Though examined in relatively few studies, transsynaptic effects from individual ICMS pulses also can modulate the firing of neurons well beyond the cortical locale of the stimulating electrode. Single pulses delivered in macaque ventral premotor cortex (PMv) modulate the firing of many M1 neurons, and vice versa, single pulses delivered in M1 modulate the firing of many PMv neurons (Kraskov et al., 2011). ICMS pulses delivered in rodent somatosensory can produce activity dependent plasticity in neurons of the rostral forelimb motor area (Averna et al., 2019). Understanding such cortico-cortical effects of ICMS pulses, in addition to advancing our knowledge of both how ICMS is perceived and how ICMS influences other unperceived brain activity (Mazurek and Schieber, 2019; Mazurek and Schieber, 2021), may have practical applications. In bidirectional brain-computer interfaces (O’Doherty et al., 2011; Flesher et al., 2021; Pandarinath and Bensmaia, 2022), for example, ICMS delivered in the primary somatosensory cortex (S1) to provide artificial tactile feedback might modulate the discharge of M1 neurons being used to decode movement intent. Such ICMS-induced modulation could reduce the performance of a decoding algorithm that uses the activity of those M1 neurons as inputs (Shelchkova et al., 2022).

We therefore tested the hypothesis that ICMS pulses delivered in macaque S1 directly modulate the firing of neurons in two cortical motor areas known to receive somatosensory input. In M1, neurons respond to both cutaneous and deep peripheral stimulation (Rosen and Asanuma, 1972; Lemon and Porter, 1976; Wong et al., 1978; Fetz et al., 1980). In PMv, neurons also respond to both cutaneous and deep peripheral stimulation, but have larger and more complex somatosensory receptive fields (Rizzolatti et al., 1981; Gentilucci et al., 1988; Rizzolatti et al., 1988; Graziano and Gandhi, 2000). We went on to examine the spatial distribution of the modulation that S1-ICMS pulses produced in each of these cortical motor areas.

## Materials and Methods

We performed new analyses on data collected during a previously reported study (Mazurek and Schieber, 2017a). Methods for behavioral training, neural recording, and delivering ICMS, all have been described there in detail and are summarized here only briefly, whereas the new experimental design and statistical analyses for the present work are described here in detail.

### Non-human Primates

Two male rhesus monkeys (Monkeys L and X, weight 9 and 11 kg, age 11 and 13 years old, respectively) were subjects in the present study. All procedures for the care and use of these non-human primates followed the Guide for the Care and Use of Laboratory Animals and were approved by the University Committee on Animal Resources at the University of Rochester, Rochester, New York.

### Initial Training with Visual Instructions

Each monkey was initially trained to perform a reach-grasp-manipulate (RGM) task as described in detail previously (Mollazadeh et al., 2011; Mazurek and Schieber, 2017a). The monkey initiated each trial by reaching to a central home object, grasping it, and pulling it for a variable initial hold period (700-1500 ms). Then the monkey was instructed to reach to, grasp, and manipulate 1 of 4 peripheral objects by illumination of a ring of blue LEDs around the base of that object. The 4 objects were arranged at 45⁰ intervals 13 cm radially from the home object. The monkey pushed a button, turned a sphere, pulled a coaxial cylinder (coax), or pulled a perpendicular cylinder (perp). The monkey then held the object in the manipulated position for a final hold period of 700-1000 ms. Three hundred milliseconds after the successful completion of a trial, a water reward was delivered, and the trial ended. A 1000 ms inter-trial interval occurred before another trial could be initiated. Trials for each target object were presented pseudo-randomly in blocks. Each block included one trial for each target object, and the order of the 4 instructed objects within a block was re-randomized between blocks. Errors occurred if the monkey failed to hold the home object for the required duration, failed to release the home object within 1000 ms of the instruction onset, contacted the wrong target object, failed to contact a peripheral object within 1000 ms of releasing the center object, or failed to hold the target object for the required duration. When an error occurred, trials were aborted immediately, no reward was given, and the same object was instructed again on the next trial to prevent the monkey from avoiding trials involving a particular peripheral object. Trials following an error were excluded from analysis because the monkey could have known which object would be instructed. The RGM task was controlled by a custom software written in TEMPO (Reflective Computing, Olympia, WA).

### Neural Recordings

Once trained to perform the RGM task instructed with visual cues, each monkey was implanted with multiple 16-electrode (impedance ∼0.5 MΩ, 70% Pt, 30% Ir) floating microelectrode arrays (FMAs, Microprobes for Life Sciences, Gaithersburg, MD) in the primary somatosensory cortex (S1), primary motor cortex (M1), and premotor cortex (PM) of the left hemisphere, as described previously (Mollazadeh et al., 2011). The present analyses are based on ICMS pulses delivered via 4 S1 arrays in each monkey while spikes were recorded from 4 M1 arrays in each monkey and from 4 PM arrays in monkey L, but from only 2 PM arrays in monkey X. In monkey L, all 4 PM FMAs were located in the ventral premotor cortex (PMv), while in monkey X no arcuate spur was present and the more medial array (D) may have been in the dorsal premotor cortex. Nevertheless, for present purposes we will refer to all of these PM arrays as having been in PMv. The intracortical locations of electrode tips were confirmed by the presence of single- or multi-unit activity. Neural data were recorded with a Multi Acquistion Processor and SortClient software (Plexon, Inc., Dallas, TX). After 1,000 – 32,000x amplification, waveforms that crossed a threshold set online by the user were sampled at 40 kHz and saved for off-line sorting along with behavioral event markers generated by the TEMPO system. After off-line sorting (Plexon Offline Sorter), the signal-to-noise ratio (SNR) and the estimated percentage of inter-spike interval (ISI) violations (< 1ms) were used to classify each sorted unit as a definite single unit (SNR ≥ 3 and no ISIs < 1ms), a probable single unit (SNR ≥ 2.5 and <10% ISIs < 1ms), a multi-unit (SNR ≥ 1.5 and <90% ISIs < 1ms), or noise (SNR < 1.5 or ≥ 90% ISIs < 1ms) (Rouse and Schieber, 2016).

### Learning to Use ICMS Instructions

Several months after the microelectrode arrays had been implanted, each monkey was trained to perform the RGM task using ICMS pulse trains delivered in S1 as instructions, instead of the visual LED cues (Mazurek and Schieber, 2017a). ICMS trains consisted of symmetric, biphasic, cathodic-leading, 200 µs per phase, 1-64 µA pulses delivered at a constant frequency. For monkey L, ICMS pulse frequency was 100 Hz for all target objects, whereas for monkey X, ICMS pulse trains had a different frequency for each object: button – 75 Hz, sphere – 100 Hz, perp – 150 Hz, coax – 225 Hz. ICMS pulses were produced by a constant-current IZ2 Neural Stimulator controlled by an RZ5 BioAmp Processor hosted by a PC running the OpenEx Software Suite (Tucker-Davis Technologies, Gainesville, FL).

Each of the 4 objects was instructed with a train of ICMS pulses delivered through electrodes on a different S1 array. ICMS pulses for each instruction were delivered simultaneously through a set of 3-7 electrodes all on the same S1 array. At first, these ICMS pulse trains were delivered concurrently with the LED cues, i.e. stimulation on the S1 array assigned to the instructed target was delivered simultaneously with illumination of the LED ring surrounding that target. Then, over multiple subsequent daily sessions, the LEDs were gradually dimmed until they could remain completely off while the monkey continued to perform well. At this point, ICMS trains served as the only instruction for the correct target object on each ICMS-instructed trial, with no visual cue. ICMS trains were delivered throughout the reaction time and movement time on each trial (∼400 — 600 ms), from instruction onset (the first ICMS pulse) until the monkey had manipulated an object. Ultimately, trials with only visual instructions or only ICMS instructions could be interleaved randomly and the monkeys each continued to perform well. The data used in the present analyses were recorded in four such sessions from monkey L and three from monkey X.

### Experimental Design and Statistical Analysis

All analyses presented here were performed off-line using MATLAB (MathWorks, Natick, MA).

#### Peristimulus Time Histograms

To determine whether M1 and PMv neurons were directly modulated by pulses of S1-ICMS, we constructed four peristimulus time histograms (PSTHs) for each recorded unit – one PSTH triggered on pulse artifacts from each of the four different ICMS instructions. The time of each ICMS pulse artifact was identified offline by discriminating the waveforms of stimulation artifacts on recording channels from M1 or PMv in which any spikes were much smaller than the recorded artifacts. Each rhythmic train of artifacts could be unambiguously assigned to ICMS pulses from a given S1 array based on the shapes of the four different artifact waveforms and knowledge of which object was instructed on each trial. The time window of each PSTH was chosen to span from 20% of the inter-pulse interval before the timestamp of the triggering ICMS pulse artifact until the timestamp of the next ICMS pulse artifact. For example, a 100 Hz pulse train would have 10 ms inter-pulse intervals, and therefore the PSTH window spanned from −2 ms before to +10 ms after the trigger time. A PSTH bin-width of 0.1 ms was chosen to provide high temporal resolution. Raw spike counts in each bin were divided by the total number of ICMS trigger pulses, such that the PSTH ordinate represented the probability of the neuron firing a spike in each 0.1 ms bin (Guggenmos et al., 2013). Figure 2 illustrates one of the four PSTHs (for instruction 1, the button) for a definite single unit in M1.

In forming these PSTHs, we blanked any neural data from −0.2 ms before to +0.6 ms after the timestamp assigned to each stimulus artifact by Offline Sorter, as we considered any spike waveform on an M1 or PMv recording channel that occurred during this interval to have been potentially distorted by collision with the ICMS pulse artifact. This produced a minimum blanked window in each PSTH from −0.2 ms to +0.6 ms around the trigger time at 0.0 ms. ICMS pulse artifacts varied across different recording channels, however, often having waveforms different from the artifacts in the channels used to trigger each PSTH. In some recording channels, the artifacts could begin more than −0.2 ms before the timestamp or last longer than +0.6 ms after the timestamp. We therefore expanded the duration of the artifact window separately for each PSTH, using the 15^th^ percentile of all the bins in each PSTH as a threshold. We defined the first bin that exceeded the 15^th^ percentile proceeding backward from time −0.2 ms to be the start and the first bin that exceeded the 15^th^ percentile proceeding forward from time +0.6 ms to be the end of the artifact window on that channel. Any spikes within this “final artifact window” (shaded pink in all PSTH figures) were excluded from subsequent analyses.

In one situation, however, we terminated the final artifact window earlier than the 15^th^ percentile threshold crossing proceeding forward from time +0.6 ms. As illustrated by the example PSTH of Fig. 3A, the artifact trigger time at 0.0 ms sometimes was followed by a prolonged series of bins with zero spike counts, suggesting that the ICMS pulses produced early inhibition of the unit. We therefore set a maximum termination time for artifact windows to distinguish early inhibition following ICMS pulses from spikes lost due to collisions with the ICMS artifacts. To determine this maximum termination time, we identified the time of the earliest spike discriminated more than the +0.6 ms minimum after the artifact trigger for each of the 596 constructed PSTHs. We then compiled a histogram of those earliest spike times (Fig. 3B), which appeared bimodal. We interpreted the first mode to represent PSTHs in which spikes occurred promptly following the ICMS pulse artifact (i.e. without early inhibition) and the second mode to represent PSTHs in which spikes resumed after a period of early inhibition. We fit a generalized Gaussian mixture model to the bimodal distribution and used the nadir between the two components at 2.3 ms after the trigger time at 0.0 ms as the maximum possible termination of an ICMS pulse artifact window. Any artifact window (determined using the 15^th^ percentile criterion) that lasted longer than 2.3 ms therefore was terminated at 2.3 ms. Whereas bins during the final artifact window were excluded from subsequent analyses, a series of zero-count bins beyond 2.3 ms was considered to indicate a period of early inhibition rather than collisions with artifacts, and therefore was included in subsequent analyses. The number of triggers used to construct PSTHs varied for a number of reasons. First, while ICMS was delivered at constant frequencies through the reaction and movement epochs, the monkeys’ reaction and movement times varied from trial to trial. When reaction and movement times were longer, more ICMS pulses were available as triggers. Second, for monkey X the instruction for each object had a different frequency (button – 75 Hz, sphere – 100 Hz, or perp – 150 Hz) and higher frequency pulse trains therefore provided more triggers. And third, ICMS artifacts occasionally were distorted by collision with other waveforms and consequently were unrecoverable in the Offline Sorter. For each object, Table 1 summarizes the medians and interquartile ranges across all trials in all sessions for each monkey’s reaction times (Rx), movement times (Mv), and response times (Rsp, reaction + movement times), as well as the number of triggers (nTrig) used to construct PSTHs.

**Table 1.** Behavioral metrics during ICMS-instructed trials determining the number of triggers available for PSTH construction. For each object and each monkey, the median, the 25^th^ percentile, and the 75^th^ percentile across all trials in all sessions is given for: Rx – reaction time, Mv – movement time, Rsp – total response time, nTrig – number of triggers.

| Behavioral Metrics |  | Button |  |  | Sphere |  |  | Perp |  |  | Coax |  |  |
| --- | --- | --- | --- | --- | --- | --- | --- | --- | --- | --- | --- | --- | --- |
|  |  | Med | 25<sup>th</sup> | 75<sup>th</sup> | Med | 25<sup>th</sup> | 75<sup>th</sup> | Med | 25<sup>th</sup> | 75<sup>th</sup> | Med | 25<sup>th</sup> | 75<sup>th</sup> |
| L | Rx | 390 | 343 | 480 | 293 | 262 | 322 | 302 | 274 | 332 | 356 | 318 | 409 |
|  | Mv | 275 | 249 | 303 | 249 | 226 | 280 | 223 | 201 | 243 | 223 | 209 | 241 |
|  | Rsp | 668 | 622 | 774 | 542 | 510 | 574 | 522 | 493 | 551 | 581 | 541 | 639 |
|  | nTrig | 12519 | 11243 | 13804 | 7906 | 5576 | 10264 | 7958 | 6742 | 9074 | 10463 | 8355 | 12773 |
| X | Rx | 597 | 429 | 974 | 581 | 520 | 685 | 466 | 417 | 531 |  |  |  |
|  | Mv | 225 | 205 | 251 | 135 | 127 | 147 | 131 | 123 | 143 |  |  |  |
|  | Rsp | 841 | 653 | 1207 | 724 | 656 | 850 | 604 | 547 | 671 |  |  |  |
|  | nTrig | 11683 | 9586 | 13620 | 8507 | 7869 | 8727 | 7726 | 6900 | 10453 |  |  |  |

#### Modulation in PSTHs

In the PSTH of each neuron-array pair, we identified excitatory peaks and/or inhibitory troughs when ≥ 3 consecutive bins went beyond ± 2 standard deviations of the baseline mean calculated from the bins preceding the final artifact window. In cases for which the mean minus 2 standard deviations were less than zero, we required 7 consecutive bins with counts of zero to identify a trough. For 59 pairs we did not attempt to identify troughs because the total spike count was less than the number of bins in the PSTH (i.e. <100 spikes for 100 Hz pulse trains leading to 100 bins, <133 spikes for 75 Hz pulse trains leading to 133 bins, etc.), producing many bins that contained zero spikes by chance alone.

We then measured the latency, full width at half maximum (FWHM), and amplitude of each peak or trough, as illustrated in Figure 4. Latencies were calculated as the time from the beginning of the final artifact window to the first bin in which a peak or trough crossed above or below 2 standard deviations from the mean of the baseline spike probability. FWHM was calculated as the difference in time between the bins in which a peak or trough crossed 50% of its maximum or minimum value with respect to the baseline mean. We quantified amplitude as the maximum of a peak or the minimum of a trough minus the baseline mean, normalized as a percentage of the baseline mean. For peaks, we also calculated the cumulative probability as the sum of the probability in each bin from onset to offset of the peak.

#### Code Accessibility

Analysis-specific code and data are available from the Lead Contact upon request.

## Results

During each session, neuron spiking data were collected from all the 16-channel microelectrode arrays implanted in M1 and PMv of each monkey. After off-line sorting, each sorted waveform was classified as a definite single unit (DSU), probable single unit (PSU), or multi-unit (MU) according to criteria described in the Methods. Units were tracked across consecutive days based on their mean waveforms, mean firing rates, autocorrelograms, and cross-correlograms with other units (Fraser and Schwartz, 2012). Units found to be present in multiple sessions were analyzed only once, from the first session in which they appeared. In Monkey L, 81 units were analyzed – 38 from M1 and 42 from PMv. The 38 M1 units were comprised of 9 DSUs, 8 PSUs, and 21 MUs. The 42 PMv units were comprised of 5 DSUs, 12 PSUs, and 25 MUs. In Monkey X, 69 units were analyzed – 36 from M1 and 33 from PMv. The 36 M1 units were comprised of 6 DSUs, 14 PSUs, and 16 MUs. The 33 PMv units were comprised of 2 DSUs, 6 PSUs, and 25 MUs. To obtain the best available estimate of the prevalence of excitatory and inhibitory effects, we chose to include not only DSUs and PSUs, but also MUs in the analyzed population. The presence of a peak or trough in the PSTH of an MU demonstrated that at least one neuron recorded at that location was modulated by the S1-ICMS pulses.

### Direct modulation of neurons in M1 and PMv produced by S1-ICMS

Figure 5 displays four PSTHs constructed with spikes from the same definite single unit recorded in M1 of Monkey L. Each PSTH was triggered on the artifacts of ICMS pulses from one of the four ICMS instructions, each delivered via electrodes on a different S1 array (colored red). Whereas the ICMS pulses of instructions 1 and 4 did not modulate this neuron’s discharge (Fig. 5A, 5D), the ICMS pulses of instructions 2 and 3 did, producing different patterns of peaks and/or troughs in their respective PSTHs (Fig. 5B, 5C). We refer to such evidence of increased and/or decreased spike probability as “direct modulation.”

ICMS instructions delivered through electrodes on the four different S1 arrays in each monkey provided 152 M1 neuron-array pairs and 168 PMv pairs in monkey L; 144 M1 pairs and 132 PMv pairs in monkey X. The pairs involving one S1 array in monkey X (colored pale yellow in Figure 1) were excluded from analysis, however, because the high stimulation frequency used for that array (225 Hz) resulted in inter-pulse intervals (4.4 ms) too short for reliable assessment of modulation. The remaining 108 M1 and 99 PMv neuron-array pairs from monkey X were included in the subsequent analyses.

**Figure 1.**
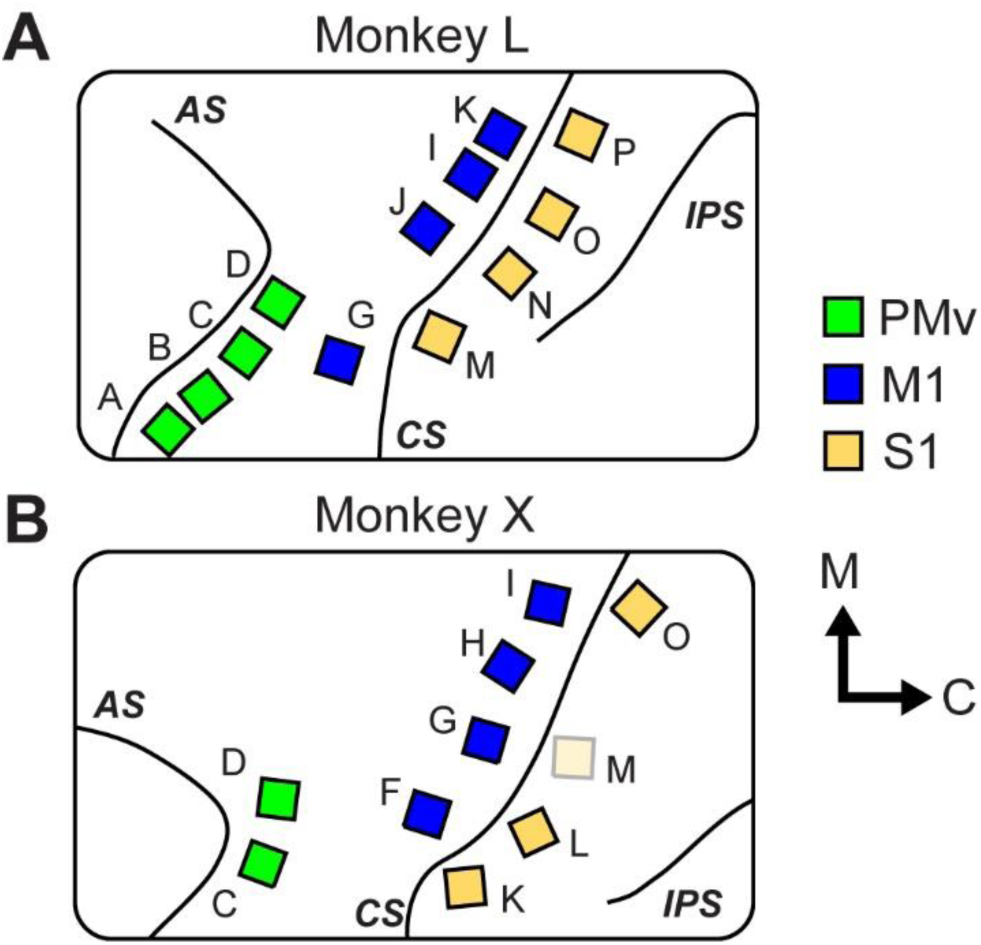
Location of microelectrode arrays implanted in Monkey L (A) and Monkey X (B). Arrays (∼2x2 mm) were implanted in primary motor cortex (M1, blue), ventral premotor cortex (PMv, green), and primary somatosensory cortex (S1, yellow). In Monkey X, data from ICMS delivered via array M (pale yellow) were not used in the present study because the high frequency (225 Hz) resulted in inter-pulse intervals (4.4 ms) too short for reliable assessment of modulation. Letters next to each array are used to identify that array. AS – arcuate sulcus, CS – central sulcus, and IPS – intraparietal sulcus. Orientation arrows indicate medial (M) and caudal (C) directions. Modified with permission from Mazurek & Schieber, 2017.

**Figure 2.**
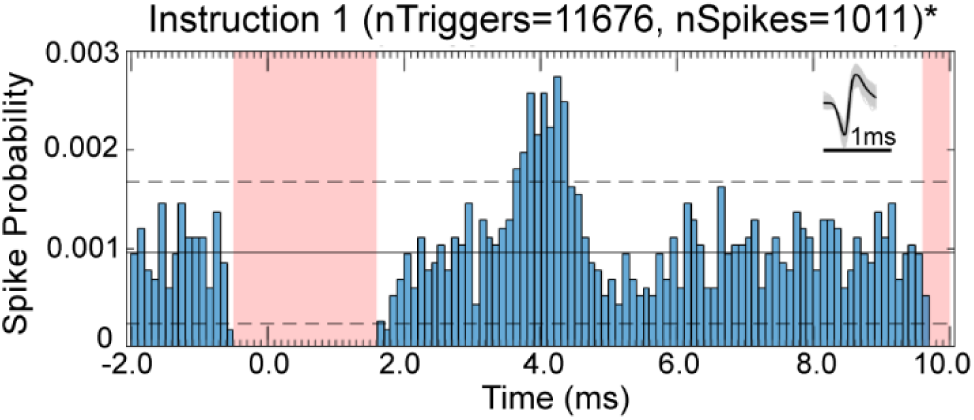
Example peristimulus time histogram. Across multiple RGM trials, this definite single unit from M1 fired a total of 1011 spikes (nSpikes) during ICMS Instruction 1 (button). Spike counts in each 0.1ms bin were converted to probabilities by dividing the count in each bin by the number of ICMS pulse artifacts used as triggers to form the PSTH (nTriggers). Pink rectangles show the artifact window during which spikes could not be reliably discriminated. Horizontal black lines represent the mean (solid line) ± 2 standard deviations (dashed lines) of the baseline bins preceding the first artifact window. The average waveform of the unit is shown in the upper right corner of the PSTH, with overlapped individual spikes in gray and a 1ms timescale bar below in black. Ch78U2, L, 20150729.

**Figure 3.**
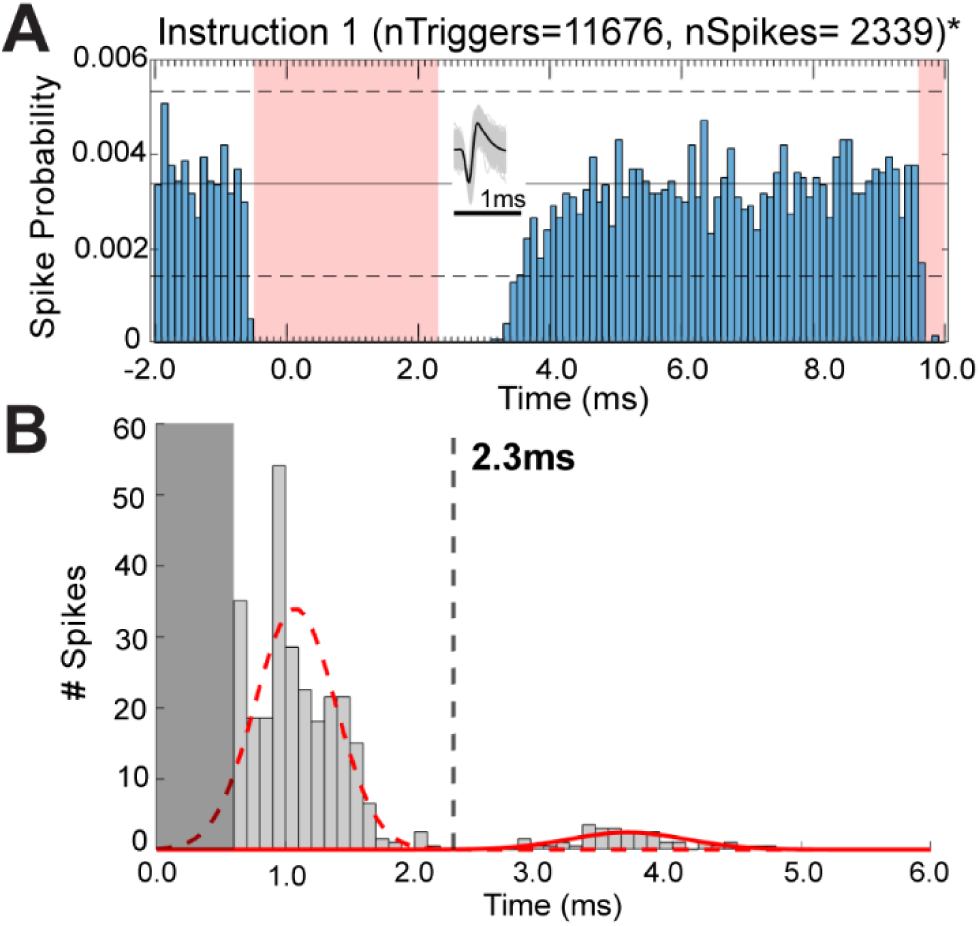
Distinguishing early inhibition from collisions with ICMS pulse artifacts. A) Example PSTH in which a period of early inhibition followed the ICMS pulse artifact. The final artifact window began −0.5 ms before and ended 2.3 ms after the trigger time at 0.0 ms. (Ch114U1, L, 20150729). B) For every PSTH, the time of the earliest spike discriminated more than the +0.6 ms minimum after the artifact trigger at 0.0 ms (indicated by the dark grey rectangle) was collected and binned in 0.1 ms steps. The resulting distribution appeared bimodal, and a generalized Gaussian mixture model with two components was fit to the data. The first component (dashed red curve) was interpreted as spikes resuming as soon as the artifact ended with no immediately following inhibition, whereas the second component (solid red curve) was interpreted as spikes resuming after an early inhibitory period immediately following the artifact. The nadir between the two components at 2.3 ms therefore was set as the maximum time following the trigger at which the final artifact window could terminate.

**Figure 4.**
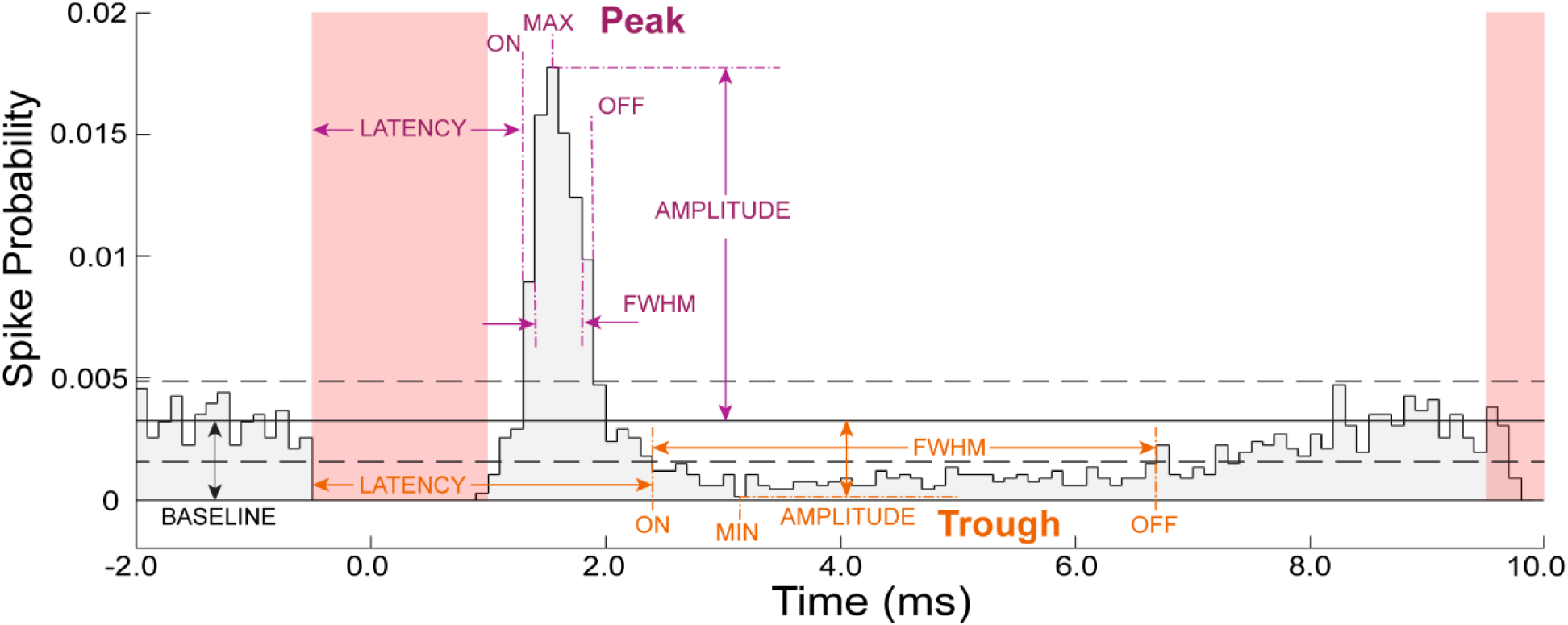
Peak and trough characteristics. Peaks and troughs were identified when the PSTH crossed 2 standard deviations (horizontal dashed lines) beyond the mean (horizontal solid line) of the baseline period before the final artifact window (pink rectangles). The onset latency, offset time, FWHM, and amplitude of each peak (purple) or trough (orange) were calculated as described in the text. In this example, the onset and offset times of the trough happened to coincide with those of the half minimum.

For M1 neurons, we identified at least one peak or trough that exceeded ± 2 standard deviations of the pre-stimulus baseline (see Methods) in the PSTHs of 85/152 (55.9%) neuron-array pairs from monkey L and 44/108 (40.7%) pairs from monkey X; for PMv neurons, 45/168 (26.8%) pairs from monkey L and 35/99 (35.4%) pairs from monkey X. We also assessed the percentage of neurons that were directly modulated by ICMS delivered through electrodes on at least one S1 array. For M1 neurons, 33/38 (87%) in monkey L and 28/36 (78%) in monkey X were directly modulated by ICMS delivered on at least one S1 array; for PMv neurons, 26/42 (62%) in monkey L and 29/33 (88%) in monkey X. The majority of neurons recorded in both M1 and PMv were directly modulated by ICMS pulses delivered on at least one S1 array.

Combining the data from the two monkeys, we identified 46 peaks in M1 and 44 in PMv; 160 troughs in M1 and 77 in PMv. The frequencies of peaks vs. troughs differed in M1 vs PMv (*Χ^2^*-test, p = 0.0052). Overall, inhibition was more common than excitation, more so in M1 than in PMv.

Although single peaks or troughs were most frequent, many PSTHs showed multiple effects, as exemplified by the two troughs in Fig. 5B or the peak and trough in Fig. 5C. Table 2 gives the numbers of PSTHs with single peaks, single troughs, or multiple effects in M1, in PMv, and in the two areas combined, each broken down by whether the unit was a DSU, PSU, or MU. We considered the possibility that MUs might be more likely to show multiple effects, but pooling the effects from M1 and PMv showed no dependence between unit quality and single peak vs. single trough vs. multiple effects (*Χ^2^*-test, p=0.088). Comparing the DSU+PSU+MU totals for M1 vs. PMv, however, did confirm that the frequencies of single peak vs. single trough vs. multiple effects differed in the two cortical areas (*Χ^2^* test, p=2.82e-05), with troughs and multiple effects being more frequent in M1 and peaks more frequent in PMv.

**Figure 5.**
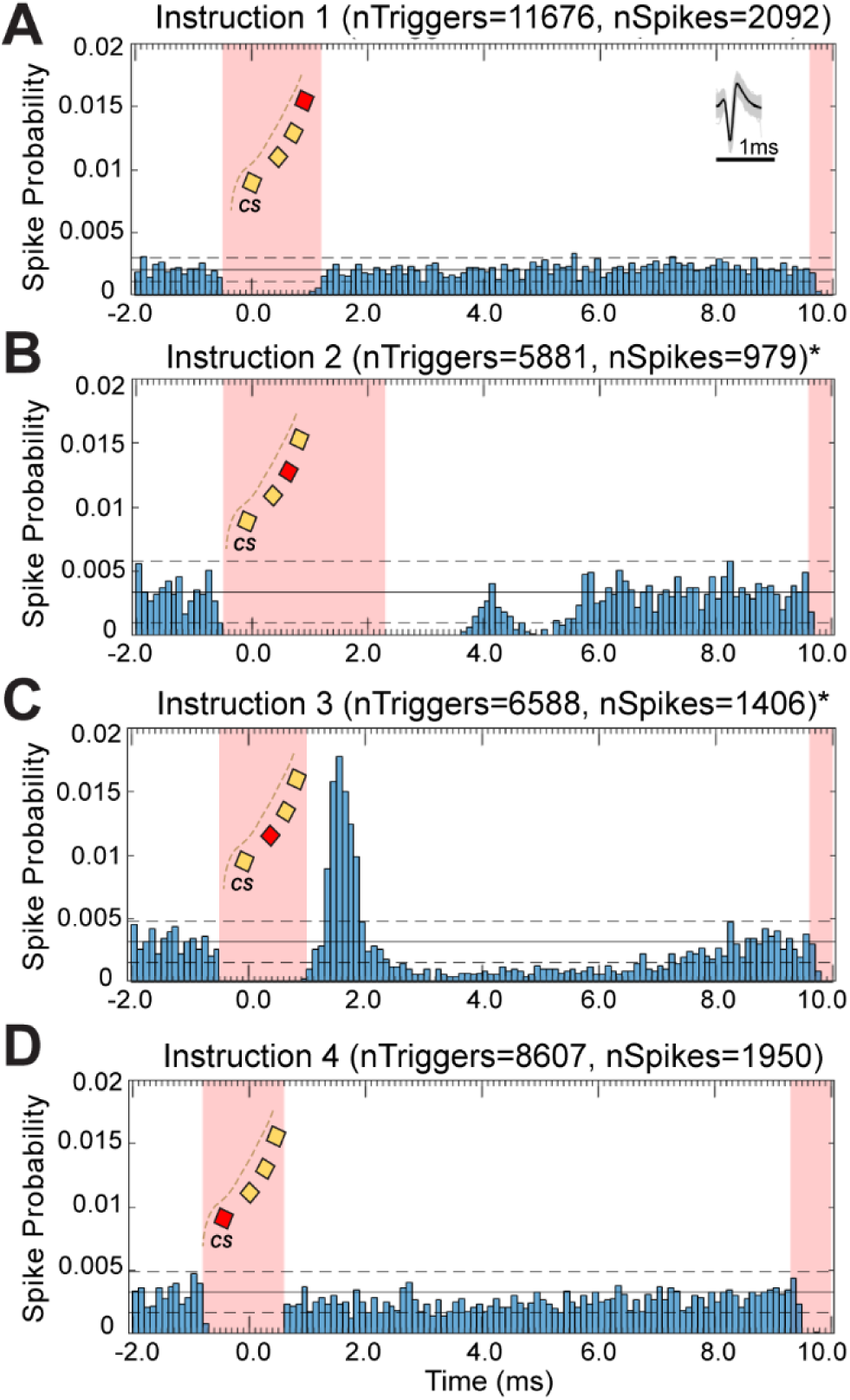
Peristimulus time histograms triggered on ICMS pulse artifacts from the instruction for each target object. All four PSTHs show spikes from the same definite single unit in M1, triggered on ICMS pulse artifacts from the A) button, B) sphere, C) perpendicular cylinder, or D) coaxial cylinder instruction. Spike counts in each 0.1 ms bin were converted to probabilities by dividing the count in each bin by the number of ICMS pulse artifacts used as triggers to form the PSTH (nTriggers). Pink rectangles show the artifact window during which spikes could not be reliably discriminated. Insets within the left pink rectangle illustrate the arrays implanted in S1 posterior to the central sulcus (CS), with the stimulated array colored red and the others yellow. Horizontal black lines represent the mean (solid line) ± 2 standard deviations (dashed lines) of the baseline bins preceding the first artifact window. The average waveform of the unit is shown in the upper right corner of the PSTH in A, with overlapped individual spikes in gray and a 1 ms timescale bar below in black. Note that the baseline firing probability varied among the four PSTHs because the neuron fired differently during trials involving the four different objects. Ch109U1, L, 20150729

**Table 2.** Types of direct modulation. M1 – primary motor cortex, PMv – premotor cortex, DSU – definite single unit, PSU – probable single unit, MU – multi-unit

| Region | Unit Quality | 1 peak | 1 trough | Multiple | Total |
| --- | --- | --- | --- | --- | --- |
| M1 | DSU | 1 | 6 | 6 | 13 |
|  | PSU | 2 | 23 | 9 | 34 |
|  | MU | 2 | 43 | 37 | 82 |
|  | **Total** | **5** | **72** | **52** | **129** |
| PMv | DSU | 3 | 0 | 1 | 4 |
|  | PSU | 6 | 9 | 4 | 19 |
|  | MU | 11 | 25 | 21 | 57 |
|  | **Total** | **20** | **34** | **26** | **80** |
| M1 + PMv | DSU | 4 | 5 | 8 | 17 |
|  | PSU | 8 | 32 | 13 | 53 |
|  | MU | 13 | 68 | 58 | 139 |
|  | **Total** | **25** | **105** | **79** | **209** |

### Latency, full-width half-maximum, and amplitude of direct modulation effects in M1 and PMv neurons

We then quantified the latency, full-width at half maximum, and amplitude of each peak or trough. Fig. 6A shows the distributions of peak (left) and trough (right) latencies in M1 (top row) and PMv (bottom row). Although the median latency of troughs was not different in M1 versus PMv (1.8 ms in M1 vs. 1.6 ms in PMv, p=0.49, Mann-Whitney U-Test), the median latency of peaks was longer in M1 than in PMv (4.3 ms in M1 vs. 3.4 ms in PMv, p=0.04, Mann-Whitney U-Test). Inspection of the distributions shows that this difference was not due to fewer short-latency peaks produced in M1, but rather due to more peaks at longer latencies (> 4ms) – an unexpected finding given that M1 is physically closer than PMv to S1.

**Figure 6.**
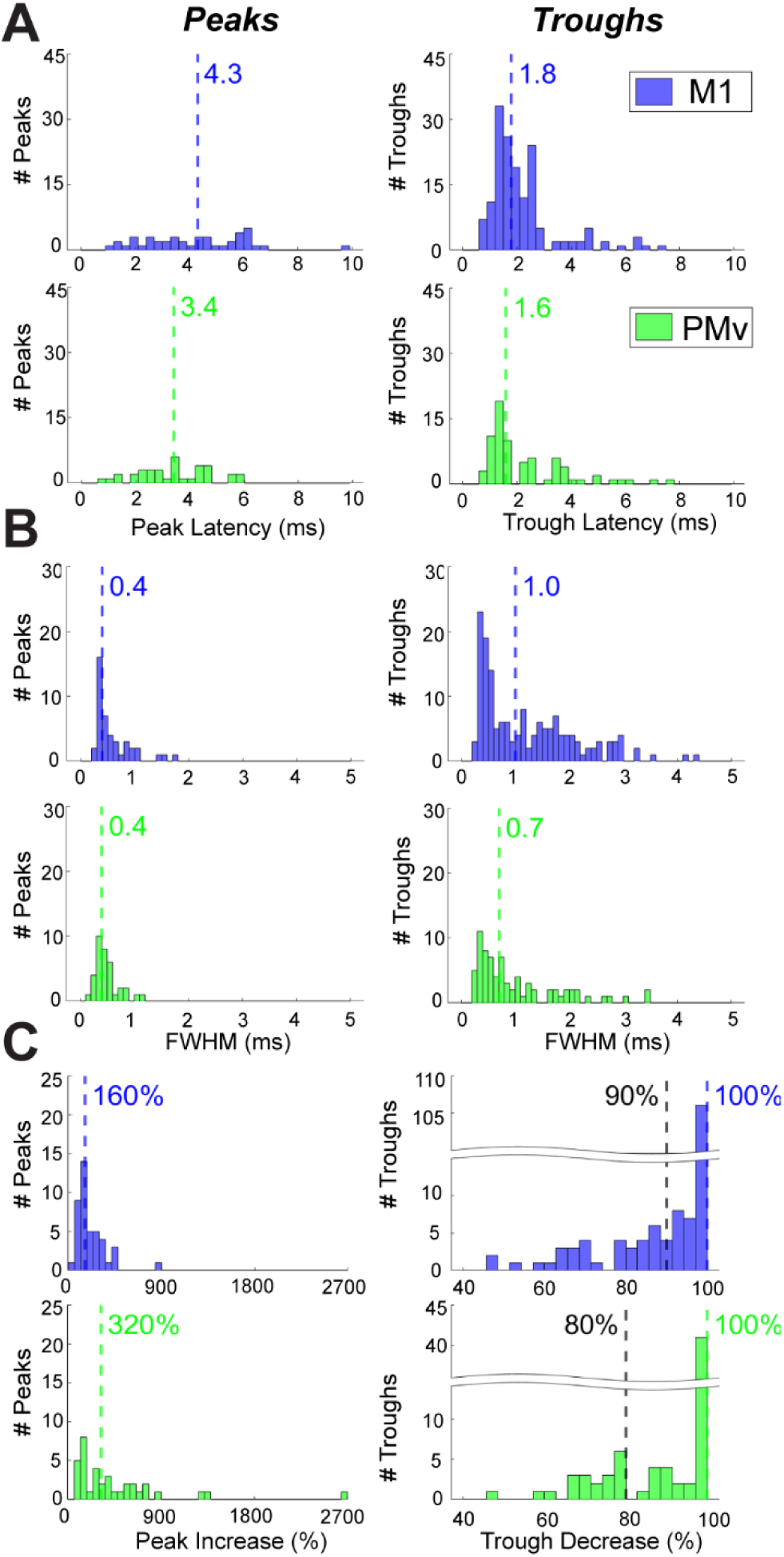
Properties of excitatory and inhibitory effects observed in M1 (blue) and PMv (green) neurons following S1-ICMS pulses. Separate histograms are shown for peaks (left column) and troughs (right column). Dashed vertical lines mark the median of each distribution. A) Latencies. The range of latencies was similar for excitation and inhibition in both M1 and PMv, Short-latency troughs were more common than short-latency peaks in both areas. Histogram bin size is 0.3 ms. B) Full-Width Half-Maxima (FWHM). Most peaks and troughs in both M1 and PMv were of relatively short duration, though some troughs lasted longer than any peaks. Histogram bin size is 0.1 ms. C) Amplitude (percent change from baseline). Black dashed lines mark medians after excluding all values of 100%, i.e. the floor for inhibition, 0 spikes. Histogram bin size is 60% for peaks and 3% for troughs.

We quantified the duration of each peak or trough by measuring its full-width at half maximum (FWHM). The distributions of FWHMs (Fig. 6B) show that virtually all peaks and many troughs in both M1 and PMv were relatively short-lived, with FWHMs < 1 ms. In contrast to peaks, though, many troughs had FWHMs > 1 ms, making the median duration of troughs longer than that of peaks in both areas (1.0 ms vs. 0.4 ms in M1, p=1.0e-06, Mann-Whitney U-Test; 0.7 ms vs 0.4 ms in PMv, p=2.8e-05, Mann-Whitney U-Test). Between M1 and PMv, no differences were found in FWHM for either peaks (p=0.34, Mann-Whitney U-test) or troughs (p=0.13, Mann-Whitney U-test).

We quantified the amplitudes of peaks/troughs as the maximal/minimal value expressed as a percent change from the baseline. Distributions are shown in Fig. 6C. Peak amplitudes were as high as 869% in one M1 neuron and 2650% in one PMv neuron (corresponding to the probability of a spike in the peak bin of 2.5e-03 and 7.5e-03, respectively). The median percent increase for peaks was significantly higher in PMv than in M1 (320% in PMv vs 160% in M1, p=0.009, Mann-Whitney U-Test).

Our ability to assess trough amplitude was constrained by the spike probability floor of zero, which was reached by most troughs, resulting in a median percent decrease of 100% for both areas. We recalculated the median percent decrease after excluding values of 100%. The median percent decrease of the remaining troughs was still substantial in both areas: 90% for M1 and 80% for PMv.

The relatively short latency of some peaks raised the possibility that S1-ICMS might have excited some neurons in M1 or PMv antidromically. Antidromic responses would be expected to produce very narrow peaks with cumulative probabilities approaching 1.0, reflecting one-to-one following. However, antidromic responses evoked by stimulation of thin, unmyelinated axon collaterals may not follow stimulation at frequencies ∼100 Hz, with as few as 10% of stimulus pulses evoking a somatic action potential, i.e. a spike probability of 0.1 (Chomiak and Hu, 2007). In Fig. 7A we have plotted the FWHM of each peak we evoked with S1-ICMS pulses against its cumulative probability. Most peaks had cumulative probabilities < 0.05. Of the 6 peaks with cumulative probabilities > 0.05, an M1 neuron had the narrowest FHWM (arrow). The PSTH with this peak is shown above in Fig. 5C. Fig. 7B illustrates the jitter of the first action potentials of this M1 unit (black) following 10 stimulus artifacts (red), indicating that excitation of this M1 neuron was orthodromic and transsynaptic. We thus infer that most, if not all, of the peaks we found represented orthodromic, transsynaptic excitation.

**Figure 7.**
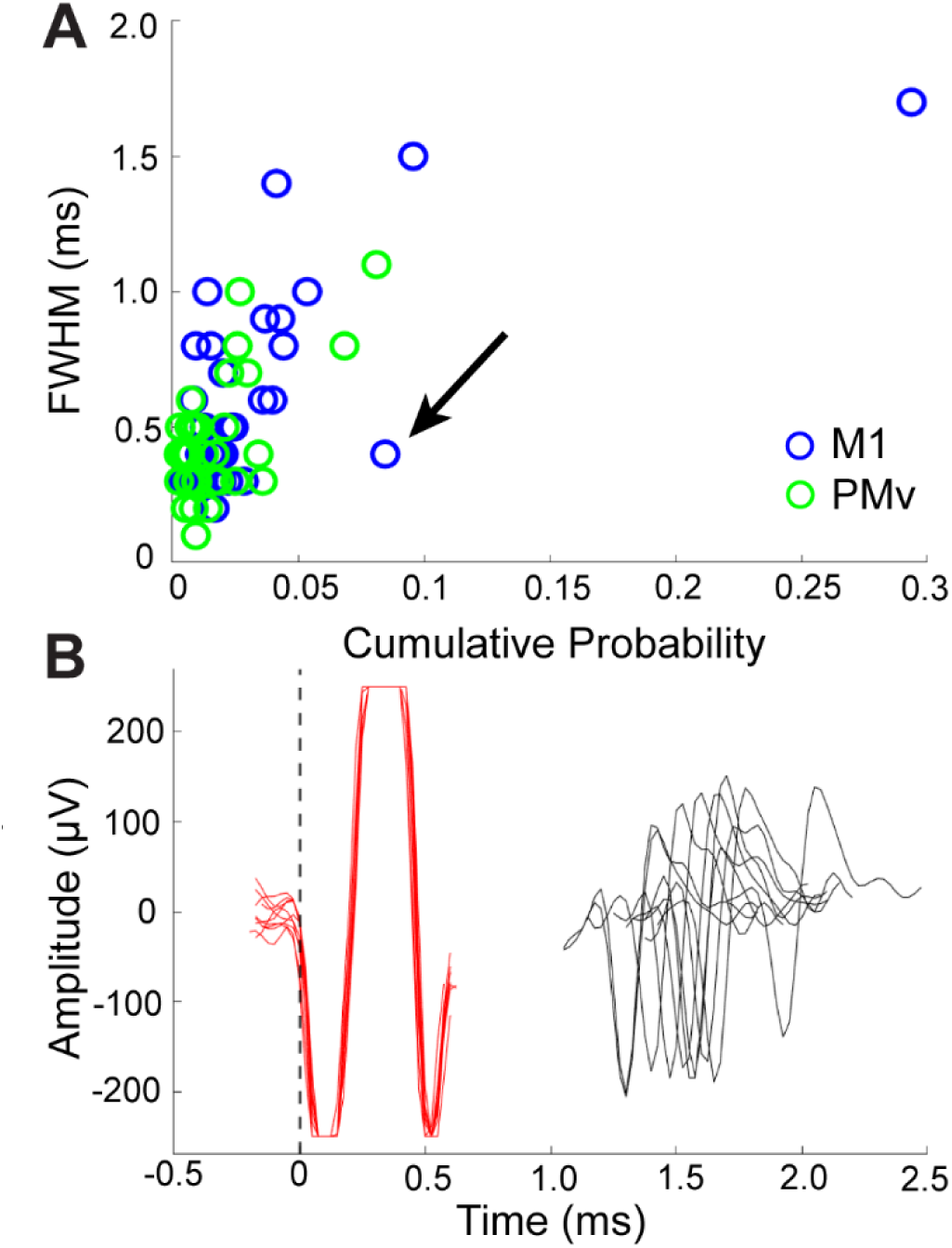
Cumulative probability vs. FWHM and Jitter of S1-ICMS effects. A) Cumulative probability versus full-width half-maximum of each peak from M1 (blue) and PMv (green). Cumulative probabilities were calculated by summing each bin in the PSTH from the onset of the peak to the offset (described in the Methods). The black arrow indicates a point that represents a peak with both a relatively high cumulative probability and a narrow FWHM, as might be expected of an antidromic response. B) Overlay of 10 ICMS pulse artifacts (red traces) and the 10 following spikes (black traces) from the neuron-array pair denoted by the black arrow in A. Both waveforms were aligned on the timestamps of the ICMS artifacts from the designated artifact channel as described in Methods (0 ms, vertical dashed line). The jitter in the latencies of the spike waveforms relative to the artifact waveforms indicates orthodromic, transsynaptic excitation of the neuron. The PSTH of this neuron-array pair is shown in Figure 5C.

### Spatial convergence and divergence of S1-ICMS effects on neurons across M1 and PMv

Next, we examined the spatial convergence of effects from multiple S1 arrays onto individual units in M1 and PMv, as well as the spatial divergence of effects from stimulation of each S1 array to units across each cortical area. We present our findings on convergence and divergence in both tabular (Fig. 8) and graphic (Fig. 9) form.

**Figure 8.**
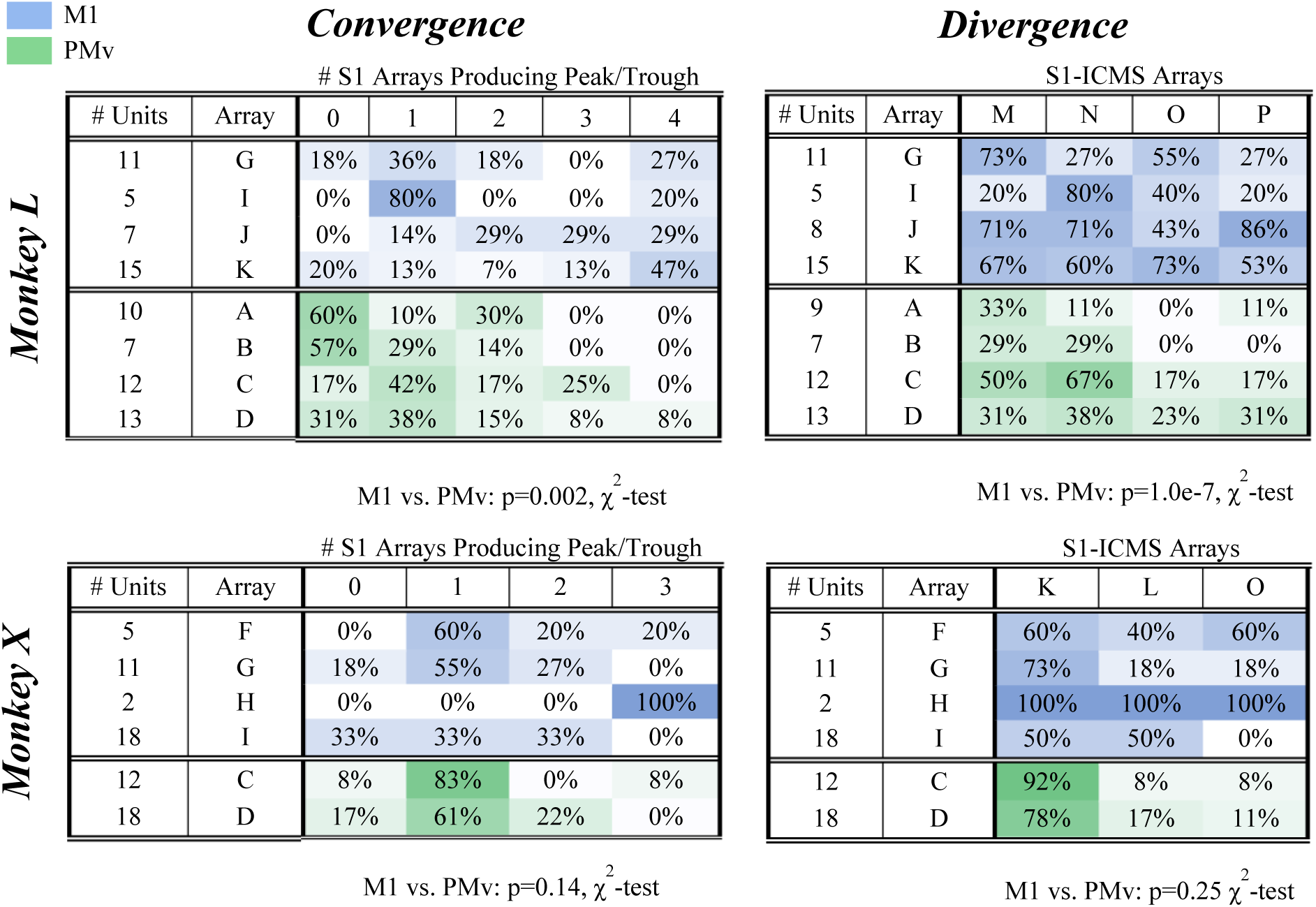
S1-ICMS effects by array. For convergence tables (left), each entry gives the percent of units on each M1 (blue) or PMv (green) array that had a significant peak and/or trough in the PSTHs triggered on artifacts of ICMS pulses delivered through different numbers of S1 arrays, from 0 to 4. For divergence tables (right), each entry gives the percent of units on each M1 or PMv array with a peak and/or trough in the PSTH triggered on artifacts from ICMS pulses delivered through the indicated S1 array. All values have been rounded to the nearest whole number and entries are color coded on a scale from white (0%) to dark blue/green (100%, M1/PMv). Results for monkey L are displayed above and results for monkey X below.

**Figure 9.**
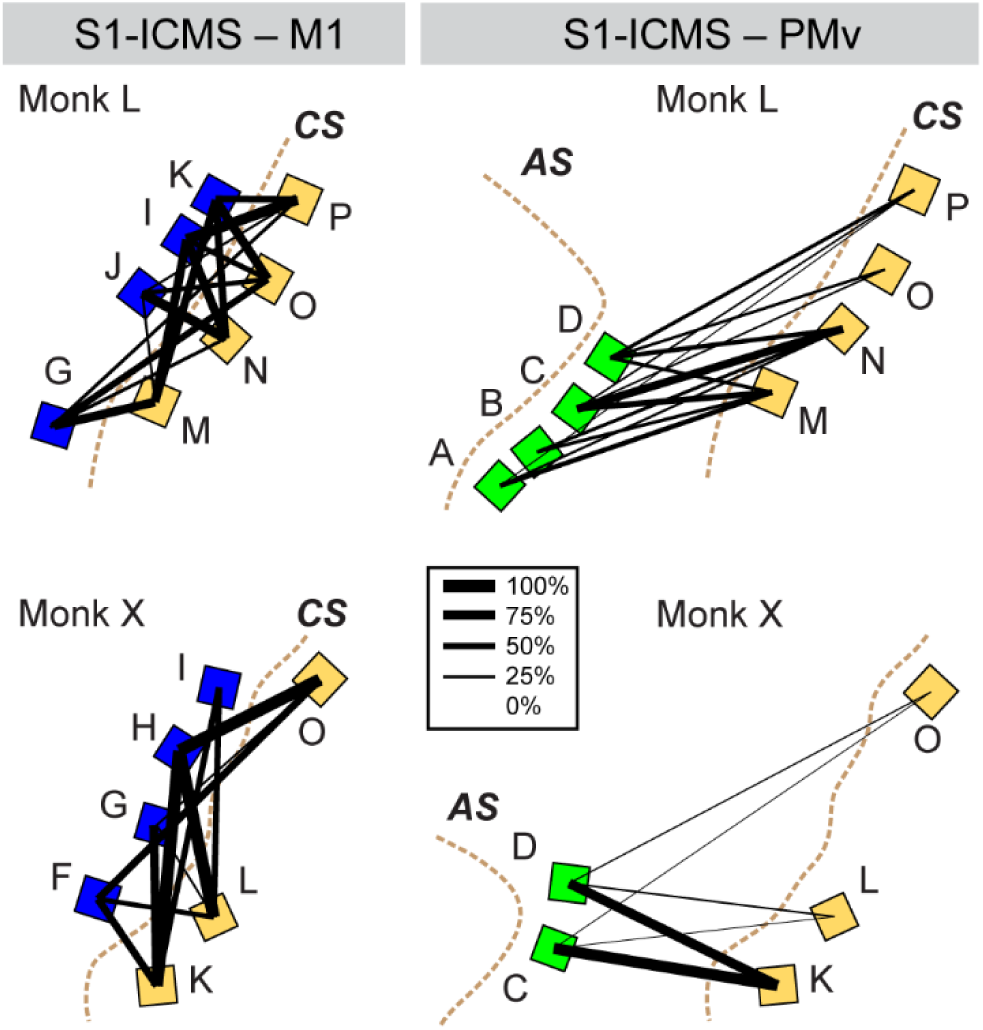
Graphic topography of direct modulation effects from each S1 array. Each square represents an FMA implanted in M1 (blue), PMv (green), or S1 (yellow). The thickness of each solid black line is proportional to the percentage of units recorded from each M1 or PMv array that showed a peak and/or trough following ICMS pulses delivered through the connected S1 array. The letters next to the arrays correspond to those in Figures 1 and 8. Brown dotted lines indicate sulci: Arcuate Sulcus (AS) and Central Sulcus (CS).

In both monkeys, ICMS effects from multiple different S1 arrays converged on individual neurons in both M1 and PMv. Figure 8 (Convergence, left) gives the percentage (rounded to the nearest whole number) of neurons recorded on each array in M1 or PMv that had at least one significant peak or trough following ICMS delivered through different numbers of arrays in S1. Focusing on the right column (4) in monkey L, for example, shows that ICMS effects converged from all four S1 arrays on some individual units recorded from each of the four M1 arrays and from one of the four PMv arrays. In monkey X, ICMS effects from converged from all three S1 arrays on individual units from two of the four M1 arrays and from one of the two PMv arrays. In total, 13/38 (34%) M1 units and 1/42 (2%) PMv units received effects converging from all four S1 arrays in monkey L, and 3/36 (8%) M1 and 1/30 (3%) PMv units received effects converging from all three S1 arrays in monkey X. In each monkey, other units received effects converging from fewer S1 arrays. ICMS effects from wide territories in S1 thus converged on many units in both M1 and PMv.

Conversely, ICMS effects from each S1 array diverged to neurons recorded on multiple different arrays in both M1 and PMv. Figure 8 (Divergence, right) gives the percentage of neurons recorded on each M1 or PMv array that had at least one peak or trough following ICMS pulses delivered through each S1 array. Looking down the columns shows that in monkey L, ICMS effects diverged to neurons on all 4 M1 arrays from each of the 4 S1 arrays and diverged to all 4 PMv arrays from 2 of the 4 S1 arrays (M and N). In monkey X, ICMS effects diverged to neurons on all 4 M1 arrays from 2 of the 3 S1 arrays (K and L) and diverged to neurons from both PMv arrays from all three S1 arrays.

We then compared the convergence and divergence of S1-ICMS effects between M1 and PMv. By comparing counts of units receiving direct modulation from 0, 1, 2, 3, or 4 S1 arrays, we found that in both monkeys individual M1 units were directly modulated by ICMS pulses delivered on more S1 arrays than were individual PMv units. This trend was significant in monkey L (p=0.002, χ^2^ test), though not in monkey X (p=0.14, χ^2^ test). We also found that in both monkeys S1-ICMS delivered on individual arrays diverged to modulate units across M1 more commonly than PMv, though again this trend was significant in monkey L (p=1.0e-7, χ^2^ test), but not in monkey X (p=0.25, χ^2^ test). As illustrated by the thickness of the black lines in Figure 9, both convergence and divergence of ICMS effects were more extensive from S1 to M1 than from S1 to PMv.

## Discussion

We found that ICMS pulses delivered in S1 modulated the spiking activity of a majority of units recorded in M1 and PMv. In both areas, we found inhibitory effects to be more common than excitatory effects. In addition, the duration of inhibitory effects was often longer than that of excitatory effects. S1 outputs thus may produce more inhibition than excitation in both M1 and PMv.

We also found extensive divergence of effects from S1 to both M1 and PMv. ICMS pulses delivered simultaneously via a few electrodes on a single ∼2x2 mm array in S1 produced effects in neurons recorded on multiple other ∼2x2 mm microelectrode arrays spread throughout the upper extremity representation in M1 and over a considerable territory in PMv. Conversely, many single-or multi-units anywhere in these regions of M1 and PMv received effects converging from all arrays spread across the S1 upper extremity representation. Although macaque S1 has a detailed somatotopic organization (Pons et al., 1985; Pons et al., 1987), its outputs to M1 and PMv appear to have less somatotopic segregation.

To our knowledge, few if any previous studies have examined the responses produced in multiple cortical areas by ICMS in another cortical area. Our simultaneous recordings in M1 and PMv during S1-ICMS permit certain comparisons of the effects evoked in these two regions. In particular, although the underlying reasons are not apparent, M1 neurons showed a larger preponderance of inhibitory troughs than PMv neurons, whereas PMv neurons showed on average larger excitatory peaks than M1 neurons (Figure 6).

### Physiological and anatomical bases of distant effects produced by S1-ICMS

Consistent with the natural sensory inputs known to reach M1 and PMv (see Introduction), we found that more than 60% of single- and multi-units in both areas were directly modulated by ICMS pulses delivered through at least one S1 microelectrode array. The pathways via which natural somatosensory input reaches M1 and PMv are not entirely understood, though a substantial fraction may be processed through S1. We were not able to obtain histological verification of the locations of our microelectrode tips, but based on the distance of our S1 arrays from the central sulcus (< 3 mm) and their lengths (1.5 – 6.0 mm), the tips of our S1 electrodes were probably located in areas 3b and 1, with some possibly in area 3a. Area 3a projects directly to M1 (area 4), but areas 3a, 3b and 1 all project to area 2, which in turn projects to area 4 (Jones et al., 1978; Pons and Kaas, 1986; Dariansmith et al., 1993; Tokuno and Tanji, 1993; Padberg et al., 2019). Area 2 also projects to area 5, which projects more heavily to area 4 (Strick and Kim, 1978; Zarzecki et al., 1978). All of these parietal areas also project to the secondary somatosensory area (S2) in the insular cortex, which in turn projects to area 4 (Friedman et al., 1986; Dariansmith et al., 1993).

Whereas multiple transsynaptic routes are available from S1 to M1, routes from S1 to PMv are more limited. Known somatosensory projections to PMv arise only from area 3a and from S2 (Godschalk et al., 1984; Matelli et al., 1986; Kurata, 1991). The greater variety of transsynaptic routes from S1 to M1 may account in part for the distribution of latencies for peaks in M1 being slightly wider, with a slightly longer median latency compared to PMv (Fig. 6A).

The latencies and durations we observed were consistent with those observed in other studies following natural and electrical stimulation of peripheral and cortical neurons. The latencies of M1 responses to natural stimulation of the upper extremity or electrical stimulation of the median nerve range from 8 to 15 ms (Lemon, 1981). As would be expected, we found the latencies of effects produced in M1 by ICMS pulses delivered in S1 (0.9 – 9.9 ms for peaks, 0.6 – 7.2 ms for troughs) to be shorter than latencies from stimulation in the periphery. Although our S1-ICMS pulses were delivered at frequencies of ∼100 Hz, providing some potential to miss long latency effects, the latencies we observed in M1 neurons with S1-ICMS were in the same range as those found in M1 neurons in a study using PMv-ICMS pulses delivered at 10 Hz (1.8 –3.0 ms for peaks, 2–5 ms for troughs) (Kraskov et al., 2011). In humans, S1-ICMS at 100 Hz produced effects in M1 neurons at similar latencies (2–6 ms) (Shelchkova et al., 2022). Moreover, the durations of the S1-ICMS effects we found in M1 (FWHMs 0.2 – 1.7 ms for peaks, 0.3 – 4.3 ms for troughs), were similar to those obtained with PMv-ICMS (durations ∼1ms for peaks, 5–7 ms for troughs) (Kraskov et al., 2011).

A surprising observation was that the range of latencies we found for excitatory peaks in PMv neurons (0.6 – 5.8 ms) was somewhat shorter than for those in M1 neurons (0.9 – 9.9 ms), even though the physical distance from S1 to PMv is longer than from M1 to S1. Though the present latencies in PMv neurons might seem unexpectedly short, they are similar to the range reported from area 5 to M1 (0.9 – 4.4 ms) (Zarzecki et al., 1978), which covers a similar physical distance. These observations may indicate that conduction velocities are faster from S1 to PMv than to M1. In addition, the longer latencies of some effects in M1 neurons from S1-ICMS may reflect the presence of more diverse transsynaptic routes from S1 to M1 than to PMv.

We consider it unlikely that the peaks we identified represented antidromic conduction. To our knowledge, there is no evidence that PMv projects directly to S1. For M1, antidromic responses would be unlikely because axons from area 4, though projecting to areas 3a and 2, do not project to areas 3b or 1 (Godschalk et al., 1984; Matelli et al., 1986; Dariansmith et al., 1993), where most of our electrode tips probably were situated. Furthermore, for the neuron-array pair having a peak with both a high cumulative spike probability and a narrow FWHM (Figure 7), the first spike following the stimulus artifact showed temporal jitter consistent with transsynaptic excitation. Therefore, the effects that S1-ICMS pulses evoked in M1 and PMv neurons in this study were most likely orthodromic.

### Implications of Distant S1-ICMS Effects

S1-ICMS produced more inhibition than excitation in both M1 and PMv. Somatosensory inputs typically have been assessed by determining the receptive fields from which neurons in S1, M1, or PMv can be excited. We therefore had not anticipated that S1-ICMS would produce both more frequent and longer duration inhibitory effects than excitatory effects in both M1 and PMv neurons. In contrast to the present results with S1-ICMS, PMv-ICMS has been found to produce more excitatory than inhibitory modulation of M1 neurons (Kraskov et al., 2011). We suggest that, while PMv generally facilitates activity in M1, S1 may provide inhibition of undesired activity in both M1 and PMv.

We also observed extensive convergence and divergence between our arrays that spanned the upper extremity representation in S1 and our arrays in both M1 and PMv. This might have been expected in PMv given the large somatosensory receptive fields of PMv neurons (Rizzolatti et al., 1981; Graziano and Gandhi, 2000). Less expected perhaps, was the similarly extensive convergence and divergence between our S1 arrays and those in M1, which spanned the M1 upper extremity representation. A recent study in humans likewise found that ICMS on many S1 electrodes evoked responses in M1 neurons, mostly consistent with orthodromic, transsynaptic responses (Shelchkova et al., 2022). Interestingly, focal injection of tracers in area 3a, 3b, 1, or 2 results in terminal labeling spread mediolaterally over several millimeters in the same cortical area (Jones et al., 1978). Likewise neurons at any focal location in the M1 upper extremity representation send horizontal collaterals throughout that representation (Huntley and Jones, 1991). Horizontal collaterals in both S1 and M1 thus may provide an anatomical substrate for the extensive convergence and divergence we observed physiologically between S1 and M1.

### Limitations of the Present Study

The data analyzed here were collected in a study that deterimined whether ICMS delivered through microelectrodes in different S1 locations produced experiences monkeys could distinguish. To maximize the likelihood that the monkeys would distinguish S1-ICMS at different locations, we delivered current pulses of 1-64 µA simultaneously on a set of 3-7 electrodes all on the same 16-electrode array. In monkey X, we additionally varied the frequency of the pulse trains on different arrays from 75 to 225 Hz, though in monkey Q all pulse trains were delivered at 100 Hz. The data analyzed here was collected after we found S1-ICMS patterns that produced experiences each monkey could distinguish and report by performing different movements. Subsequent testing showed that the monkeys could reliably detect S1-ICMS on some single electrodes with currents as low as 10-15 µA, at frequencies as low as 30-50 Hz, or in trains as short as 400 ms. Hence the present results likely approach the upper bounds of the strength and spatial distribution of direct modulation in M1 and PM neurons produced by multichannel S1-ICMS. Stimulating simultaneously on fewer electrodes and/or at lower currents could be expected to produce fewer and weaker effects, though not necessarily with narrower spatial distribution.

Furthermore, we focused here on S1-ICMS because ICMS in S1 currently is being developed to provide somatosensory feedback for bidirectional brain-computer interfaces. We plan subsequent studies of the effects evoked by ICMS in other cortical areas as well. In addition, future studies will be needed to address questions such as i) whether the amplitude of direct modulation produced by individual ICMS pulses varies depending on task-related variation in neuron firing rates, ii) whether the use of ICMS versus visual cues as instructions for different movements affects the average firing rates of M1 and PMv neurons during task performance, and iii) how the instructional information provided by ICMS propagates across cortical areas.

## Conclusions

Whereas previous studies typically have delivered ICMS through a single microelectrode, the present results are based on ICMS pulses delivered simultaneously through multiple microelectrodes on the same ∼2x2 mm array. We chose to deliver ICMS in this manner to make it more likely that our monkeys would perceive some component of the multi-electrode S1-ICMS and thereby be able to use it as an instruction (Mazurek and Schieber, 2017a). Interestingly, recent studies of macaque and human S1 have suggested that stimulating simultaneously through multiple microelectrodes may be a valuable approach for evoking artificial sensations. Multi-channel S1-ICMS evokes tactile sensations more focal (Greenspon et al., 2023b), at shorter latencies (Sombeck and Miller, 2019; Bjanes et al., 2022), and with more distinguishable levels of perceived force (Greenspon et al., 2023a) than stimulating through a single microelectrode. Multi-electrode ICMS therefore may be used increasingly in bidirectional brain-computer interfaces (O’Doherty et al., 2011; Flesher et al., 2021). Our results show that such multi-electrode S1-ICMS can directly modulate a majority M1 and PMv neurons, with extensive spatial divergence and convergence across large cortical territories. In bidirectional BCIs that use S1-ICMS to deliver artificial feedback, these effects will need to be taken into account when decoding the intended movement from neural activity in M1 and PMv.

## Conflict of interest

The authors declare no conflicts of interest.

## Acknowledgements

The authors thank Gil Rivlis for custom task-control and stimulation control software, and Marsha Hayles for editorial comments. This work was supported by grants F31NS129099 (BR), F32NS093709 (KAM), and R01NS107271 (MHS) from the National Institute of Neurological Disorders and Stroke. The funders had no role in study design, data collection or interpretation, nor in the decision to submit the work for publication. KAM is currently a Data Scientist in the Neurology Artificial Intelligence Program, Department of Neurology, Mayo Clinic, Rochester, MN, 55905

## Author contributions

BR, KAM and MHS designed the study. KAM collected and processed the data and performed some preliminary analyses. BR performed the analyses and wrote the manuscript. MHS supervised, provided resources, and edited the manuscript.

## Notes

### Competing Interest Statement

The authors have declared no competing interest.

### Summary of Updates

Counts in text changed to match numbers reported by Figure 8; Minor changes to Figure 1 and text for clarification purposes; Added Limitations section

